# The structure of a novel ferredoxin – FhuF, a ferric-siderophore reductase from E. coli K-12 with a novel 2Fe-2S cluster coordination

**DOI:** 10.1101/2023.07.04.547673

**Authors:** I.B. Trindade, F. Rollo, S. Todorovic, T. Catarino, E. Moe, P.M. Matias, M. Piccioli, R.O. Louro

**Affiliations:** Instituto de Tecnologia Química e Biológica António Xavier da Universidade Nova de Lisboa, Avenida da República (EAN), 2780-157 Oeiras, Portugal; Departamento de Química, Faculdade de Ciências e Tecnologia, Universidade Nova de Lisboa, 2829-516, Caparica, Portugal; iBET – Instituto de Biologia Experimental e Tecnológica, Apartado 12, 2780-901 Oeiras, Portugal; Magnetic Resonance Center and Department of Chemistry, University of Florence, Via L. Sacconi 6 50019 Sesto Fiorentino, Italy

## Abstract

Iron is a vital element for life. However, after the Great Oxidation Event, the bioavailability of this element became limited. To overcome iron shortage and to scavenge this essential nutrient, microorganisms use siderophores, secondary metabolites that have some of the highest affinities for ferric iron. The crucial step of iron release from these compounds to be subsequently integrated into cellular components is mediated by Siderophore-Interacting Proteins (SIPs) or Ferric-siderophore reductases (FSRs).

In this work, we report the structure of an FSR for the first time. FhuF from laboratory strain Escherichia coli K-12 is the archetypical FSR, known for its atypical 2Fe-2S cluster with the binding motif C-C-X_10_-C-X_2_-C. The 1.9 Å resolution crystallographic structure of FhuF shows it to be the only 2Fe-2S protein known to date with two consecutive cysteines binding different Fe atoms. This novel coordination provides a rationale for the unusual spectroscopic properties of FhuF. Furthermore, FhuF shows an impressive ability to reduce hydroxamate-type siderophores at very high rates when compared to flavin-based SIPs, but like SIPs it appears to use the redox-Bohr effect to achieve catalytic efficiency.

Overall, this work closes the knowledge gap regarding the structural properties of ferric-siderophore reductases and simultaneously opens the door for further understanding of the diverse mechanistic abilities of these proteins in the siderophore recycling pathway.

## Introduction

The high abundance of reduced sulfur and ferrous iron in the Archean ocean enabled the assembly of these two elements into clusters by the emergent Life and their use as redox centers [1,2]. Today, more oxidizing conditions following the Great Oxygenation Event (GOE) have led to the precipitation of iron as ferric minerals, dramatically lowering its bioavailability [3]. Regardless, proteins containing these clusters are found ubiquitously throughout all domains of life, playing roles in essential metabolic processes such as photosynthesis and respiration. Iron-sulfur proteins comprise a family of proteins with remarkable diversity in both cofactor configuration and reduction potential extending over a 1 V range [2]. Even among proteins containing the same cluster type, reduction potentials can be modulated by the cluster environment. Furthermore, the reduction potentials of individual Fe atoms in a 2Fe-2S cluster can be differentiated to yield localized valence in odd electron states [4]. The remarkably wide range of reduction potentials and diverse structural motifs allow iron-sulfur proteins to act as electron carriers in a variety of biological processes. Ironically, this includes their function in iron acquisition pathways as the catalytic center of Ferric-Siderophore Reductases (FSRs) [5–8]. Siderophores are secondary metabolites specialized in iron scavenging [9]. However, once inside the cell, iron release from these compounds does not occur spontaneously. Instead, this process is mediated by several enzymes [10,11]. These enzymes contain either flavins as redox cofactors and are named Siderophore-Interacting Proteins (SIPs), or 2Fe-2S clusters and are designated Ferric Siderophore Reductases (FSRs) [11–13]. Generally, these enzymes promote iron release from Ferric-siderophores through the reduction of ferric iron, Fe(III), to its ferrous form, Fe(II), catalyzing the dissociation of the Fe(III)-siderophore complex. Specifically, the FSR family is characterized by the atypical cysteine binding motif C-C-X_10_-C-X_2_-C that coordinates a 2Fe-2S cluster, as found in FhuF from Escherichia coli K-12, the archetypical FSR [5]. This protein displays several unusual spectroscopic properties: i) an EPR spectrum in the reduced state with a g_z_ value of 1.994, which is smaller than that typically found in other proteins containing 2Fe-2S clusters; ii) Mössbauer spectra with a quadrupole splitting for the oxidized protein of ΔE_q_= 0.474 mm/s which is lower than that typically found in 2Fe-2S clusters, whereas for the reduced cluster the Mössbauer parameters for the ferric iron change considerably, with the quadrupole splitting increasing to 3.3 mm/s; iii) NMR spectra with unusual chemical shifts, especially in the reduced state where paramagnetic signals are found significantly downfield shifted compared to those previously reported for other proteins containing 2Fe-2S clusters [5,7,8]. To understand how the unusual structural features of FhuF that are revealed by its unprecedented spectroscopic properties govern its physiological function, here we report, the three-dimensional structure of this protein. This first structure of a member of the Ferric-Siderophore Reductase family of proteins.

The structure of FhuF reveals a different cluster coordination mode provided by the atypical cysteine-binding motif. Furthermore we provide fine details about the 2Fe-2S cluster obtained by Resonance Raman spectroscopy and additional evidence that FhuF is a ferric-siderophore reductase of hydroxamate-type siderophores.

## Methods

### Protein Expression and Purification

FhuF WT was expressed and purified as previously reported [5,7]. The mutated FhuF (FhuF-C143S) was expressed and purified as previously reported with minor modifications [5,7]. Briefly, the plasmid pKF143S that codes for His-tagged FhuF (FhuF-C143S) mutant protein was transformed into BL21 DE3 competent cells for expression. Transformed cells were grown in Luria Bertani medium supplemented with 100 mg/L ampicillin at 37 °C, 160 rpm until they reached an OD of 0.7; the temperature was then decreased to 30 °C, and cells were harvested by centrifugation after 18 hours and stored at –80 °C. The cells were later defrosted and resuspended in 20 mM Potassium Phosphate buffer pH 7.6, 300 mM NaCl with a protease-inhibitor cocktail (Roche) and DNase I (Sigma) prior to a three-pass cell disruption at 6.9 MPa using a French press. The lysate was ultracentrifuged at 204709 g for 90 min at 4 °C to remove cell membranes and debris. FhuF was then purified from the supernatant using a His-trap affinity column (GE Healthcare) where the fraction containing FhuF was eluted at 20 mM Potassium Phosphate pH 7.6, 300 mM NaCl with 250 mM imidazole. Eluted fractions were analyzed by SDS-PAGE with Blue-Safe staining (NZYTech) and UV-visible spectroscopy to select fractions containing pure FhuF. The imidazole was removed and FhuF was concentrated at 36 °C using an Amicon® Ultra Centrifugal Filter (Millipore) with a cut-off of 30 kDa. Several aliquots were flash frozen in liquid nitrogen and stored at –80 °C while others were kept at 30 °C for a week, to promote autolysis, before a second His-trap purification step was performed. The final purified FhuF (His-tag free) was concentrated from the flow-through of the second passage through the His-trap column using an Amicon® Ultra Centrifugal Filter (Millipore) with a 30 kDa cut-off. Sample aliquots were sent for N-terminus sequencing to confirm the identity and the first amino-acid residues of the purified protein samples. Aliquots were flash-frozen in liquid nitrogen for storage, or kept under anaerobic conditions for further use.

### Structure determination

N-terminally truncated and mutated FhuF (FhuF−C143S) at a concentration of 50 mg/ml was crystallized in 0.2 M Magnesium chloride hexahydrate, 0.1 M Tris-HCl pH 8.2 and 30% PEG w/v 4000 using the hanging drop vapor diffusion technique (2 μl protein:2 μl reservoir) [5,7]. Crystals were harvested and soaked in a 2 μl drop of cryo-protectant solution (0.2 M magnesium chloride hexahydrate, 0.1 M Tris-HCl pH 8.2, 30 % PEG w/v 4000 supplemented with 5 % glycerol) prior to flash-freezing in liquid nitrogen. Diffraction data were collected at 100 K to 1.9 Å resolution at ALBA Synchrotron Beamline XALOC (Barcelona, Spain). The images were processed with autoPROC, which makes use of XDS and the CCP4 suite for integration and conversion of integrated intensities to structure factors, and an anisotropic resolution cut-off was applied with STARANISO [14–18]. The data collection and processing statistics are listed in Table 1. The structure was solved by molecular replacement (MR) using PHASER in the CCP4 suite [16]. The phasing model was obtained with AlphaFold2 (through Google Colab) using the sequence of the mature FhuF protein from which the first 16 residues (Met-1 until Thr-16) were deleted [19]. The structure was rebuilt with BUCCANEER, corrected, and completed with COOT and an initial refinement was undertaken using REFMAC5 in the CCP4 suite [20–22]. Structure refinement was performed using PHENIX [23]. Throughout the refinement, the model was periodically checked and corrected with COOT against σ_A_-weighted 2|F_o_|-|F_c_| and |F_o_|-|F_c_| electron-density maps. Solvent molecules were added manually by inspection of electron-density maps in COOT. Hydrogen atoms were included in calculated positions with the PHENIX READYSET tool and isotropic displacement parameters (ADPs) were refined for all non-hydrogen atoms. In the final refinement cycles, the relative X-ray/stereochemistry and X-ray/ADP weights were optimized to reduce the gap between R-cryst and R-free. The final values of R-cryst and R-free were 21.1 % and 24.5 % respectively, with a maximum likelihood estimate of the overall coordinate error of 0.26 Å [23]. The refinement statistics are presented in Table 2. The model stereochemical quality was analyzed with MOLPROBITY and there are no outliers in the Ramachandran ϕ,φ plot [24]. The coordinates and experimental structure factors have been submitted to the Protein Data Bank with accession code 7QP5. Images were produced using PyMOL [25,26].

**Table 1.**
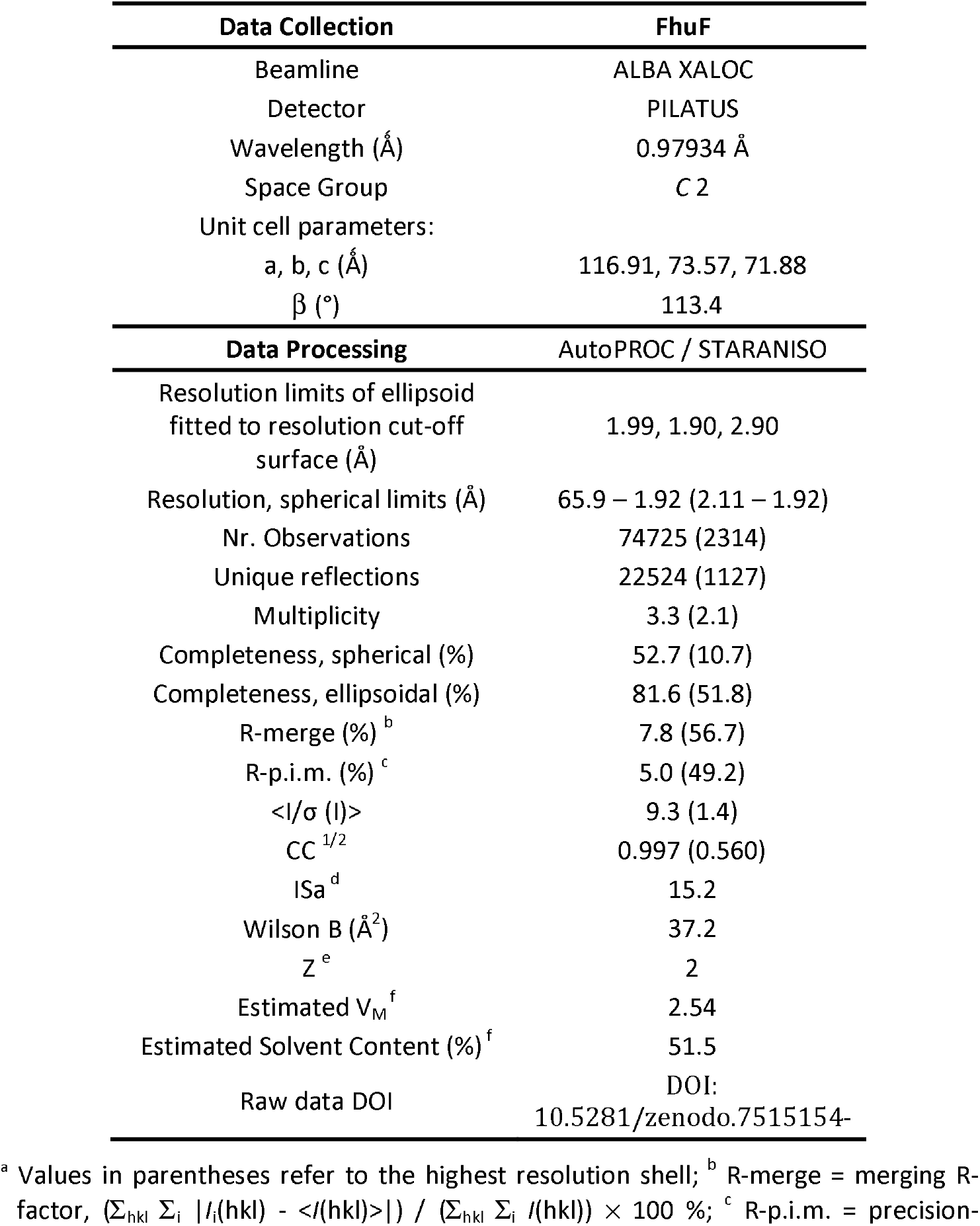
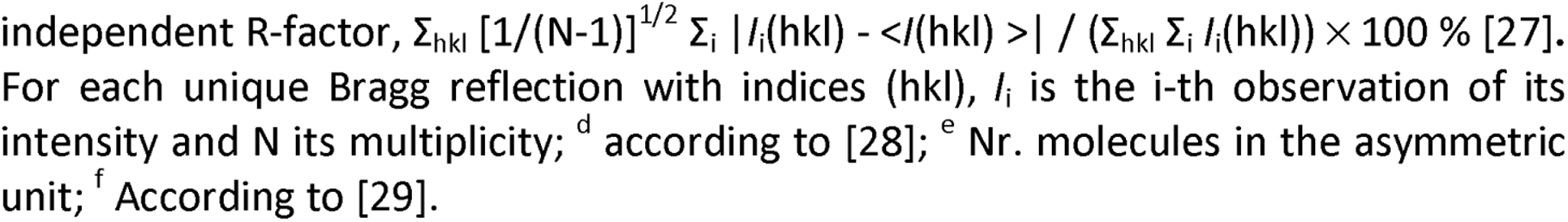
Data collection and processing statistics.

| Data Collection | FhuF |
| --- | --- |
| Beamline | ALBA XALOC |
| Detector | PILATUS |
| Wavelength (Å) | 0.97934 Å |
| Space Group | C 2 |
| Unit cell parameters: |  |
| a, b, c (Å) | 116.91, 73.57, 71.88 |
| $\beta$ (°) | 113.4 |
| Data Processing | AutoPROC / STARANISO |
| Resolution limits of ellipsoid fitted to resolution cut-off surface (Å) | 1.99, 1.90, 2.90 |
| Resolution, spherical limits (Å) | 65.9 – 1.92 (2.11 – 1.92) |
| Nr. Observations | 74725 (2314) |
| Unique reflections | 22524 (1127) |
| Multiplicity | 3.3 (2.1) |
| Completeness, spherical (%) | 52.7 (10.7) |
| Completeness, ellipsoidal (%) | 81.6 (51.8) |
| R-merge (%) <sup>b</sup> | 7.8 (56.7) |
| R-p.i.m. (%) <sup>c</sup> | 5.0 (49.2) |
| $\langle I/\sigma(I) \rangle$ | 9.3 (1.4) |
| CC <sup>1/2</sup> | 0.997 (0.560) |
| ISa <sup>d</sup> | 15.2 |
| Wilson B (Å<sup>2</sup>) | 37.2 |
| Z <sup>e</sup> | 2 |
| Estimated V<sub>M</sub> <sup>f</sup> | 2.54 |
| Estimated Solvent Content (%) <sup>f</sup> | 51.5 |
| Raw data DOI | DOI:<br>10.5281/zenodo.7515154- |

<sup>a</sup> Values in parentheses refer to the highest resolution shell; <sup>b</sup> R-merge = merging R-factor, $(\sum_{hkl} \sum_i |I_i(hkl) - \langle I(hkl) \rangle|) / (\sum_{hkl} \sum_i I_i(hkl)) \times 100$ %; <sup>c</sup> R-p.i.m. = precision-
independent R-factor, $\sum_{hkl} [1/(N-1)]^{1/2} \sum_i |I_i(hkl) - \langle I(hkl) \rangle| / (\sum_{hkl} \sum_i I_i(hkl)) \times 100 \%$ [27]. For each unique Bragg reflection with indices (hkl), $I_i$ is the i-th observation of its intensity and N its multiplicity; <sup>d</sup> according to [28]; <sup>e</sup> Nr. molecules in the asymmetric unit; <sup>f</sup> According to [29].

**Table 2.** Final refinement statistics of FhuF.

| Dataset | FhuF |
| --- | --- |
| Resolution limits (Å)<sup>a</sup> | 65.9 – 1.92 (2.00 – 1.92) |
| % $R_{\text{work}}$<sup>b</sup> | 20.9 (30.2) |
| % $R_{\text{free}}$<sup>c</sup> | 25.2 (37.9) |
| ML coordinate error estimate (Å)<sup>d</sup> | 0.26 |
| *Model composition and completeness* |  |
| Regions omitted<sup>e</sup> | 17A-18A, 17B-18B |
| Non-hydrogen protein atoms<sup>f</sup> | 3879 |
| 2Fe-2S clusters | 2 |
| Solvent molecules | 157 |
| *Mean B values (Å<sup>2</sup>)<sup>g</sup>* |  |
| Protein | 33.8 |
| 2Fe-2S clusters | 27.6 |
| Solvent | 34.6 |
| *Model r.m.s. deviations from ideality* |  |
| Bond lengths (Å) | 0.003 |
| Bond angles (°) | 0.559 |
| Chiral centers (Å<sup>3</sup>) | 0.039 |
| Planar groups (Å) | 0.005 |
| *Model validation<sup>h</sup>* |  |
| % Ramachandran outliers | 0 |
| % Ramachandran favored | 97.5 |
| % Rotamer outliers | 0.0 |
| % C<sup>β</sup> outliers | 0 |
| Clash score | 2.1 |
| PDB Accession code | 7QP5 |

<sup>a</sup> Values in parentheses refer to the highest resolution shell; <sup>b</sup> $R_{\text{work}} = (\sum_{\text{hkl}} ||F_{\text{obs}}(\text{hkl})| - |F_{\text{calc}}(\text{hkl})||) / (\sum_{\text{hkl}} |F_{\text{obs}}(\text{hkl})|) \times 100$ %; <sup>c</sup> $R_{\text{free}}$ is calculated as above from a random sample containing 5% of the total number of independent reflections measured; <sup>d</sup> Maximum-likelihood estimate by PHENIX; <sup>e</sup> Chains A and B correspond to the two independent molecules in the asymmetric unit; <sup>f</sup> Including atoms in the alternate conformations of disordered groups of residues; <sup>g</sup> Calculated from isotropic or equivalent isotropic B-values; <sup>h</sup> Calculated with MolProbity.

### Resonance Raman experiments

For resonance Raman (RR) spectroscopic experiments, 2 μL of the sample (350 µM FhuF WT in 20 mM Potassium Phosphate buffer pH 8 with 300 mM NaCl) were introduced, under strictly anaerobic conditions, into a liquid-nitrogen-cooled cryostat (Linkam), mounted on a microscope stage and cooled down to 77 K. Spectra from the frozen sample were collected in backscattering geometry using a confocal microscope coupled to a Raman spectrometer (Jobin-Yvon LabRam 800 HR, HORIBA) equipped with a 1200 mm ^-1^ grating and a liquid-nitrogen-cooled CCD detector. The 458 nm line from an Argon ion laser (Coherent Innova) was used as the excitation source. Typically, eight spectra accumulated for 60 s each using a laser power of 5.3 mW at the sample, were co-added to improve the signal-to-noise ratio (S/N). The background scattering was removed by subtraction of a polynomial function and the spectra were deconvoluted using LabSpec 5.4 software (Horiba).

### Stopped-flow experiments

The kinetic experiments were performed with HI-TECH Scientific Stopped-flow equipment (SF-61DX2) installed inside an anaerobic glove box (Mbraun MB150-GI). The temperature of the drive syringes and mixing chamber was maintained at 20 °C using a water bath. Sample solutions were prepared with 20 mM Potassium Phosphate buffer pH 8 with 300 mM NaCl and in the presence of an O_2_ scavenging system (30 mM glucose, 375 nM glucose oxidase, and 750 nM catalase). The time course of the reactions was monitored using a photodiode array. Solutions were prepared inside the anaerobic chamber with degassed water and all experiments were performed in duplicate. Data were analyzed using Kinetics Studio version 2.32 (TgK Scientific).

Ferric-siderophore reduction experiments were performed at 20 °C in the stopped-flow apparatus, using 15 μM FhuF_red_ against either 50 μM of bisucaberin or 15 μM of ferrichrome (concentrations after mixing). FhuF_red_ was prepared using sodium dithionite and then its excess was removed through buffer exchange in a HiTrap® Desalting Column (GE Healthcare). The different concentrations were used to avoid and deal with the spectra overlap between FhuF and the Fe(III)-siderophores. The reduction rate constants were obtained by fitting the kinetic traces at 452 nm with a single exponential function.

## Results and Discussion

### FhuF, an entirely novel structure

FhuF was first isolated in 1998. However, only dilute samples were stable for a few days at 4 °C, and thus the instability of FhuF and its homologs has prevented the structure determination of representatives of the FSR protein family [5,13]. The instability of FhuF was mainly translated into two main observables: first, non-reversible protein aggregation upon concentration higher than 10 mg/ml and second, protein autolysis losing the first 17 amino-acid residues in the N-terminus together with the histidine tag. Therefore, the 43 residues from the initial starting expression sequence (Supplementary Table S1) are gradually lost during the process of protein sample preparation. To increase sample concentration and sample homogeneity for crystallization experiments while simultaneously preventing protein denaturation, FhuF was kept at high concentrations at 30 °C for a week to promote faster autolysis, followed by a second His-trap purification step leaving the untagged protein in the flow-through. These two steps in sample preparation increased the sample homogeneity, and together with the use of the FhuF-C143S mutant instead of WT FhuF, decreased protein aggregation by preventing intermolecular disulfide bond formation, thus improving crystal quality and diffraction limits. The crystal structure of FhuF was determined to 1.9 Å resolution and deposited in the Protein Data Bank with accession number 7QP5 (Figure 1 and Table 2).

**Figure 1.**
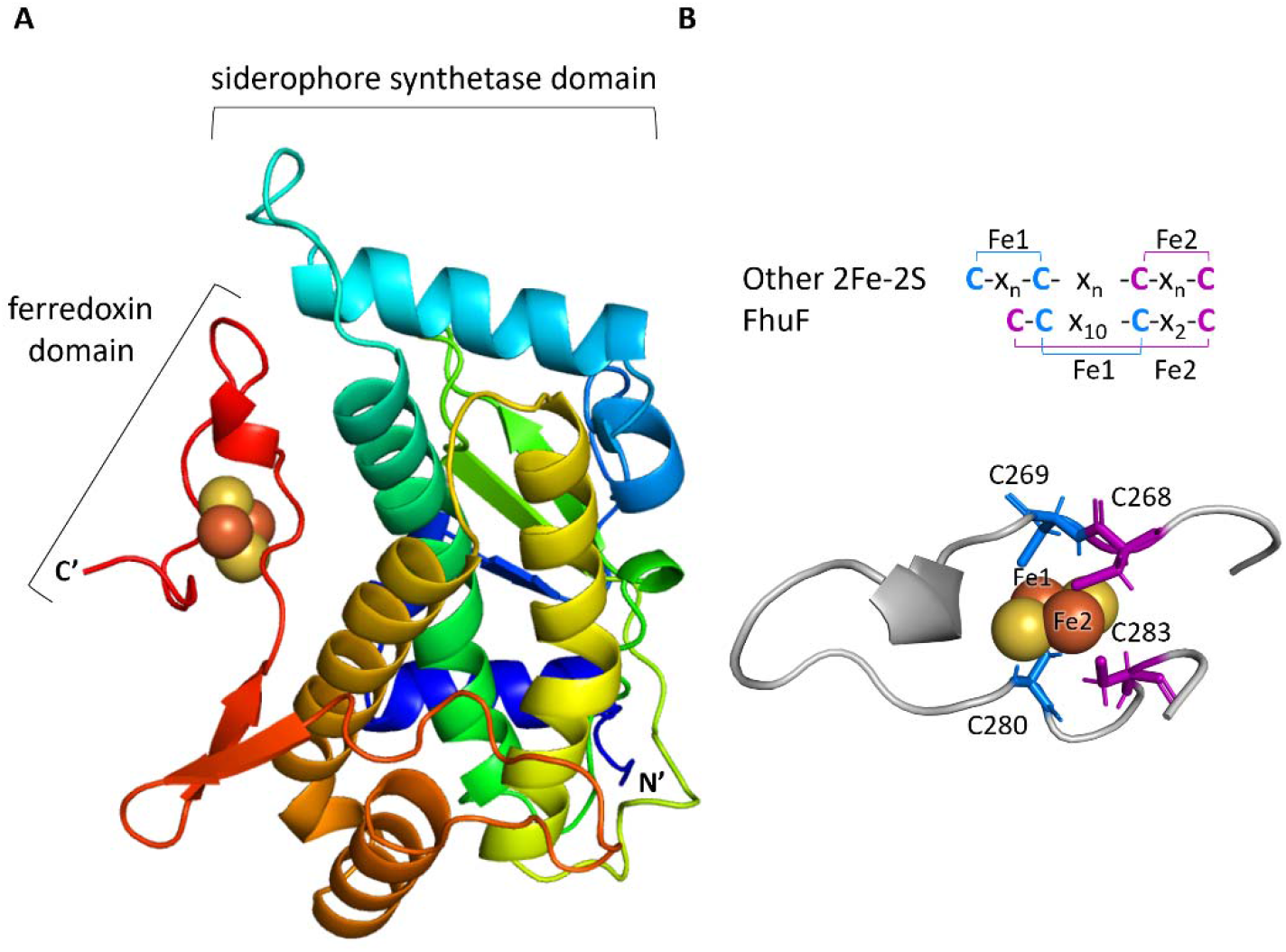
A) Cartoon diagram of the FhuF X-ray structure colored from blue to red in the N-terminal to the C-terminal direction. The atoms in the 2Fe-2S centre are represented as spheres. B) Highlight of the “ferredoxin domain” and schematic representation of the 2Fe-2S coordination mode in FhuF in comparison with other 2Fe-2S proteins where n equals 2 or more amino acid residues between cysteines.

The amino acid numbering used in this manuscript and in the deposited PDB coordinates is based on the amino acid sequence encoded by the expressed gene and thus includes 24 additional residues relative to the gene-derived protein sequence.

The structure (Figure 1 A) of FhuF reveals two domains, a 2Fe-2S cluster binding domain or “ferredoxin domain”, and a “siderophore-synthetase domain” highly similar with the “palm domain” in siderophore synthetases. This structure shows a number of remarkable features. One is the “siderophore synthetase” domain, which is unprecedented since the other family of ferric-siderophore reductases, SIPs, from the Siderophore-Interacting Proteins family, resemble oxidoreductases instead [30]. However, this agrees with the previously obtained Rosetta-derived structural model [7]. This domain is composed of a three-stranded antiparallel β-sheet that is sandwiched between a four and two α-helix bundles.

Second, the “ferredoxin” domain reveals a novel fold, where all the coordinating cysteines are within the same loop. This fold differs from those reported in all other previously characterized 2Fe-2S cluster containing proteins. The most remarkable finding is that the consensus sequence of FhuF gives rise to a novel cysteine coordination mode. Indeed, the typical coordination mode for proteins containing 2Fe-2S clusters displays two consecutive cysteines (Cys-X_2_-Cys or Cys-X-Cys) that bind the same Fe atom. Instead, in FhuF, two consecutive cysteines, that atypically also happen to be sequential neighbors, C268 and C269, bind different Fe atoms (Figure 1 B) pairing in an unprecedented way. Thus, C269 and C280 pair together and bind Fe1 (Figure 1 B) and C268 and C283 pair together to bind Fe2. Hydrogen bonds to the cluster coordinating cysteines (Figure 2 A) were also found between Cys-269 (backbone O) and Ser-117 (Sidechain H^γ^) and Asn-210 (Sidechain H^δ1^), Cys-269 (Sidechain S^γ^) with Arg-271 (backbone N-H), and Cys-268 (backbone O and N-H) with a water molecule, and then between Cys-280 (backbone O) with Cys-283 (backbone N-H). This allows us to speculate that the hydrogen bond between the backbone N-H of Arg-271 and Sγ of Cys-269 is likely responsible for the extra paramagnetically-shifted resonance found with Curie-type temperature dependence in NMR experiments [7]. The Curie temperature dependence of this signal indicates that it is associated with the ferric iron. The chemical shift of this signal is consistent with this assignment based on the distances measured in the crystal structure, using the value calculated by the hybrid density function developed for the case of single iron(III) rubredoxin [31]. This proposal provides a rationale for the steep slope observed for the temperature dependence of this signal that reflects the gradual disruption of the bond at higher temperatures.

**Figure 2.**
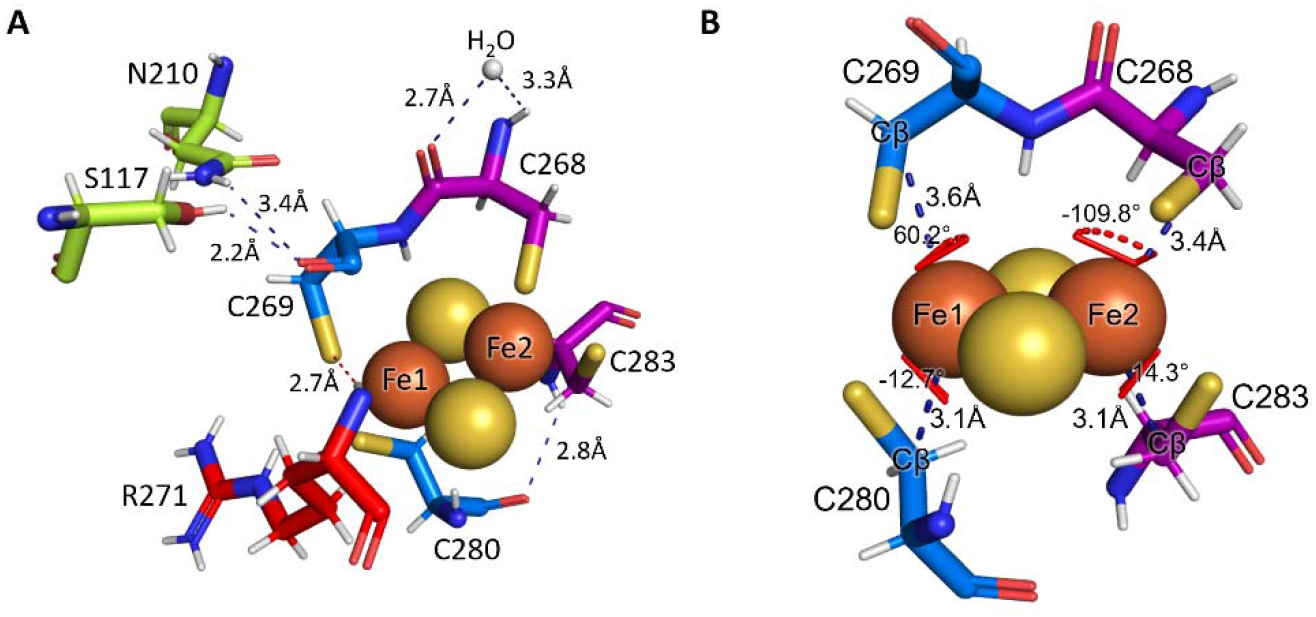
A) Diagram showing the hydrogen bonds of the cluster-coordinating cysteine residues. B) Distance and geometry of Fe-coordinating cysteines. Atom colors are dark orange (iron), yellow (sulfur). The protein residues are represented as sticks and the 2Fe-2S cluster atoms are drawn as spheres.

### The geometry of the 2Fe-2S cluster positions FhuF into the ferrochelatase class

The concept of rack-induced bonding of metal clusters in proteins to tune their reactivity was proposed by Lumry and Eyring nearly 70 years ago and used by Bo Malmström in the mid-nineteen-sixties to interpret the properties of blue copper proteins [32]. It was found that the distorted coordination imposed by the rigid organization of the protein ligands was at the origin of the unusual spectroscopic properties of the metal center. The different coordination mode of the 2Fe-2S cluster in FhuF, which lowers its symmetry vs. that of typical 2Fe-2S clusters, may be the basis for the unusual spectroscopic properties displayed by this protein [8]. This is in line with Resonance Raman (RR) spectrum of FhuF (Figure 3).

**Figure 3.**
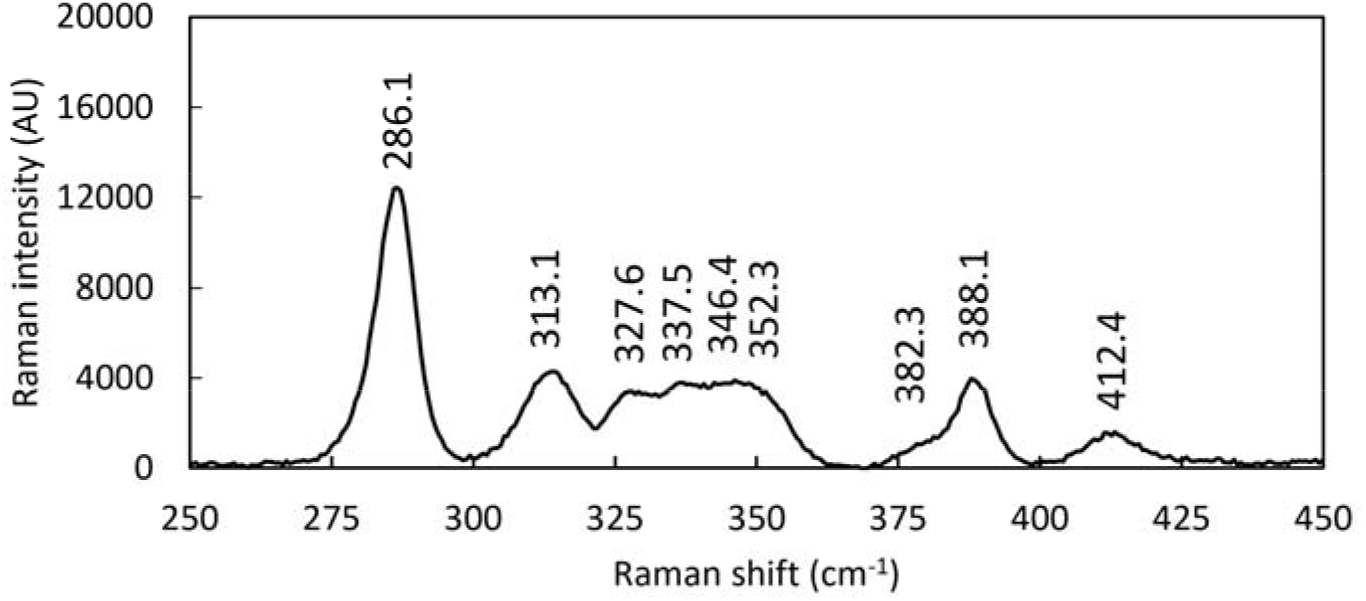
Resonance Raman spectrum of FhuF measured with 458 nm excitation, 60 s accumulation time and 5.3 mW laser power, at 77K.

RR spectra of Fe–S cluster containing proteins obtained with a laser that is in resonance with the energy of the S → Fe CT transition selectively enhance modes involving the metal–ligand stretching coordinates. RR spectra are thus sensitive to cluster type, and the modes involving predominantly bridging (Fe–S) and terminal (Fe–S) vibrations (involving inorganic sulfur and cysteinyl sulfur ligands, respectively) can be distinguished in the spectra.

The presence of ≤ 300 cm^-1^ (B_3u_) mode, which in the RR spectra of oxidized FhuF is found at 286 cm^-1^ is an unambiguous indicator of a [2Fe-2S] cluster with complete cysteinyl coordination. However, the number of identified RR bands in the spectra (Figure 3) suggests a lower symmetry and a more distorted geometry than observed in previously characterized clusters [33,34].

Additionally, the electronic structure of 2Fe-2S clusters was found to be correlated with the orientation of the ligands, which is characterized by the dihedral angle Fe-Fe-S-Cβ of the coordinating cysteines [35]. Based on the value of this dihedral angle, proteins containing 2Fe-2S clusters for which the structure is known, fall into three classes: class A congregating bona fide ferredoxins, including the thioredoxin-like ferredoxin; class B represented by xanthine oxidases; and a third class (C) for which the sole known structure to date is that of ferrochelatase [36]. Measurement of the dihedral angles Fe-Fe-S-Cβ in FhuF places it together with human ferrochelatase (PDB 1HRK) (Figure 4), considering that from an electronic point of view the sign for the dihedral angle can be exchanged [36]. This structural similarity between FhuF and ferrochelatase is also in agreement with their similar values for quadrupole splitting of the ferric iron in the Mössbauer spectra and highlights the good correlation between coordination geometry and the Mössbauer properties [8].

**Figure 4.**
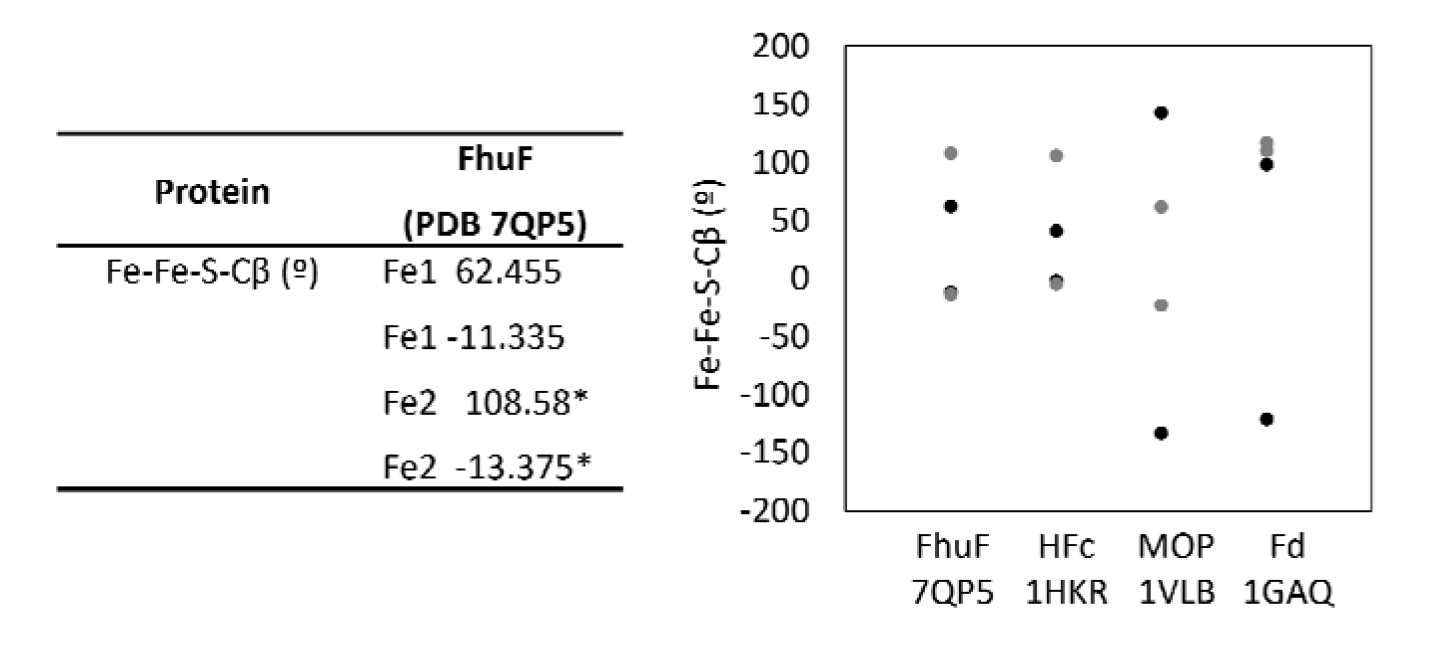
Dihedral angles Fe_i_-Fe_j_-S-Cβ for FhuF and representative members of each class of 2Fe-2S proteins according to their geometry. Class C is represented by human ferrochelatase (HFc, PDB 1HKR), class B is represented by aldehyde oxidoreductase (MOP, PDB 1VLB) and class A is represented by maize leaf ferredoxin (Fd, PDB 1GAQ). Measurements were made with PyMOL. Black circles represent angles measured with respect to Fe_j_=Fe1 and gray circles represent angles measured with respect to Fe_J_=Fe2.

### Reduced FhuF localizes the additional electron in iron 2

The structure of FhuF reported here allows us to make the connection between the valence trapping observed spectroscopically and the actual location of the additional electron in the 2Fe-2S cluster upon reduction. The EPR spectrum of reduced FhuF is distinctly rhombic even though the value of g_z_ is lower than 2, a situation unprecedented for 2Fe-2S clusters. However, coupling between a ferric and a ferrous iron can bring the value of g_z_ well below 2, in agreement with the observation of an electronic exchange coupling of 300 cm^-1^ [7,35]. This value was determined from the temperature dependence of the paramagnetically shifted NMR signals of the cysteines coordinating the cluster and is larger than the values previously reported for plant and algal 2Fe-2S ferredoxins, approaching that of Rieske proteins [37,38].

The NMR spectra of 2Fe-2S ferredoxins fall into two broad categories: one includes those obtained from plants and photosynthetic bacteria such as Anabaena, whereas the other includes those obtained from vertebrates and bacteria such as E. coli. The differences were assigned to differences in the g-tensor anisotropy of the two categories, which is rhombic for the photosynthetic ferredoxins and axial for the others [39]. As seen in the previous section, FhuF is assigned to a geometric class that is different from ferredoxins and for which there are no reported NMR spectra. The NMR spectra of FhuF do not support the classification together with ferredoxins. Indded, although the oxidized spectrum shows downshifted signals similarly to vertebrate ferredoxins, the reduced spectrum is unique and displays signals paramagnetically shifted downfield up to 200 ppm [7]. This indicates that a strong hyperfine coupling exists between the unpaired electron in the cluster and the β-CH_2_ protons of one of the coordinating cysteines.

Observation of the iron to β-CH_2_ distances for coordinating cysteines in 2Fe-2S clusters of the diverse classes of proteins shows that FhuF is the one with the greatest bond length asymmetry, with distances of 3.6 and 3.1 Å for cysteines 269 and 280, respectively. This suggests that the iron atom bound by these cysteines is the one that remains oxidized upon reduction, and that the extra electron most likely localizes in the Fe2 that is coordinated by Cys-268 and Cys-283 (Figure 2 B). In agreement with this proposal, the hydrogen bond between the backbone N-H of Arg-271 and the S^γ^ of Cys-269 (Figure 2 A) that is likely responsible for the extra paramagnetically-shifted resonance found with Curie-type temperature dependence, shows that Fe1 coordinated by Cys-269 is the ferric iron in the reduced FhuF [7]. The Fe2 coordinated by the cysteine pair Cys-268 and Cys-283 is also the Fe atom that is more solvent-exposed (Figure 6 C), further supporting the hypothesis that this is where electron localization takes place, assuming that no significant conformational changes occur upon partner interaction [4,40].

### Aspartate 259 is responsible for the redox-Bohr effect in FhuF

It was previously shown that FhuF displays redox-Bohr effect leading to a pH dependence of the reduction potential in the physiological range [7]. Unlike the Flavin co-factors of SIPs, the 2Fe-2S cluster does not perform proton-coupled electron transfer that would give rise to this pH dependence of the reduction potentials. The cysteines of the primary coordination sphere of the cluster are also not capable of undergoing acid-base transitions. The secondary coordination sphere of the cluster is known to be capable of modulating its redox potential, and the structure shows only one likely candidate for this role, the Asp 259 residue. The side chain of this amino-acid residue is found at 5.2 Å from the edge of the cluster. At this close distance, the electrostatic interaction energy between this protonatable side chain and the cluster is approximately 180 meV. This gives rise to a redox linked pK_a_ shift of approximately 3 pH units, compatible with the experimentally observed pH dependence [7].

### FhuF reduces Fe(III)-hydroxamate siderophores

It was previously hypothesized that FhuF is involved in the siderophore pathway, possibly playing a role as a ferric-siderophore reductase. This was based on the observed reduced growth of E.coli FhuF deletion mutants on plates with ferrioxamine B as sole iron source and also from observing significantly lower removal of iron from coprogen, ferrichrome and ferrioxamine B [5]. EPR experiments using reduced FhuF also showed a direct reduction of ferrioxamine E. Additionally, it was later observed that FhuF can bind both apo– and Fe(III)-rich ferrichrome [7]. Here we show that FhuF also reduces other hydroxamate siderophores, including ferrichrome and bisucaberin (Figure 5 and Figure S1).

**Figure 5.**
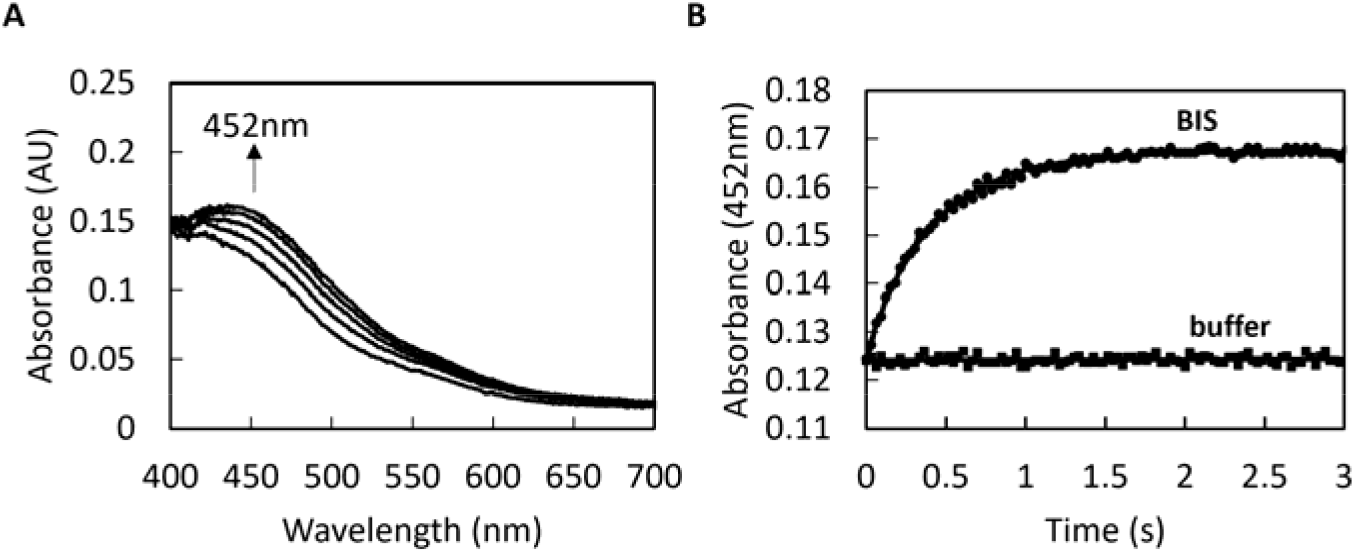
Reduction of Fe(III)-siderophores by FhuF_red_. A) Representative UV-visible spectral changes upon mixing reduced FhuF with Fe(III)-bisucaberin. B) Kinetic trace of FhuF oxidation (452 nm) upon Fe(III)-siderophore addition (BIS-bisucaberin). Data were collected at 20 °C.

All experiments were performed anaerobically in the presence of an oxygen-scavenging system since FhuF is very sensitive to oxygen even at vestigial concentrations. Despite our efforts, the reduction of FhuF was only observed with sodium dithionite (Figure S1 A) and sodium borohydride. No reduction was observed with NADH, NADPH or ferredoxin (spinach, 2Fe-2S) (data not shown), common electron donors of SIPs, the other family of ferric-siderophore reductases.

Upon mixing reduced FhuF (FhuF_red_) with Fe(III)-siderophores (bisucaberin or ferrichrome), immediate spectral changes characterized by an increase in absorption at 452 nm were observed (Figure 5 A and Figure S1 B). These are consistent with the oxidation of FhuF and reduction of Fe(III)-siderophore. Experiments were performed at 20 °C to decrease the reaction rate so that it could be measurable with the available equipment. The rate constants of Fe(III)-siderophore reduction were determined (2.74±0.08 s^-1^ for bisucaberin and 110±7.3 s^-1^ for ferrichrome) using the kinetic trace at 452 nm, which is the wavelength of maximum absorbance for a protein containing a 2Fe-2S cluster. Even though the concentrations used in this work are slightly different from those used in the work on SIP from S. frigidimarina, we demonstrate that FhuF reduces Fe(III)-siderophores faster, given the observed magnitude of difference in rate constants. This is likely because the reduction potential of FhuF is significantly lower than SfSIP (FhuF –420 mV, SfSIP –320 mV at pH≈8) thus providing a more favorable driving force for the reaction [7,30].

### The electrostatics of FhuF reveals common pockets among ferric-siderophore reductases

A visualization of the electrostatic surface potential of FhuF revealed three main charged regions. These include two positively charged regions that surround the 2Fe-2S cluster located in a neutral region between the two (pocket 1 and 2, Figure 6). Pocket 2 matches one of the regions of lower density probability found in the SAXS-derived low-resolution structure. and SwissDock predicts ferrichrome binding in this region. Furthermore, a comparison of the electrostatic surface potential of FhuF with the representatives of the SIP family (Figure 6B) [7,41,42], reveals the presence of a positively charged pocket near the catalytic center in both cases. In the SIP family, siderophores are predicted to bind in a region comprising a lysine triad. However, despite the electrostatic similarities of the pocket, this lysine triad is not found in FhuF [12,30]. It is likely that these positively charged cavities create a favorable environment for transiently accommodating Fe(III)-siderophores and drive their reduction. It has been previously observed that the redox-Bohr effect is a common feature of these enzymes [7,30]. Positively charged pockets provide an environment compatible with redox-Bohr effect. Upon electron transfer from the protein to the siderophore, the positively charged pocket favors the coupled release of a proton to maintain charge neutrality. The proton-coupled electron transfer has the mechanistic advantage of increasing the very low reduction potential of siderophores, easing their reduction, and also enhancing ferrous iron solubility.

**Figure 6.**
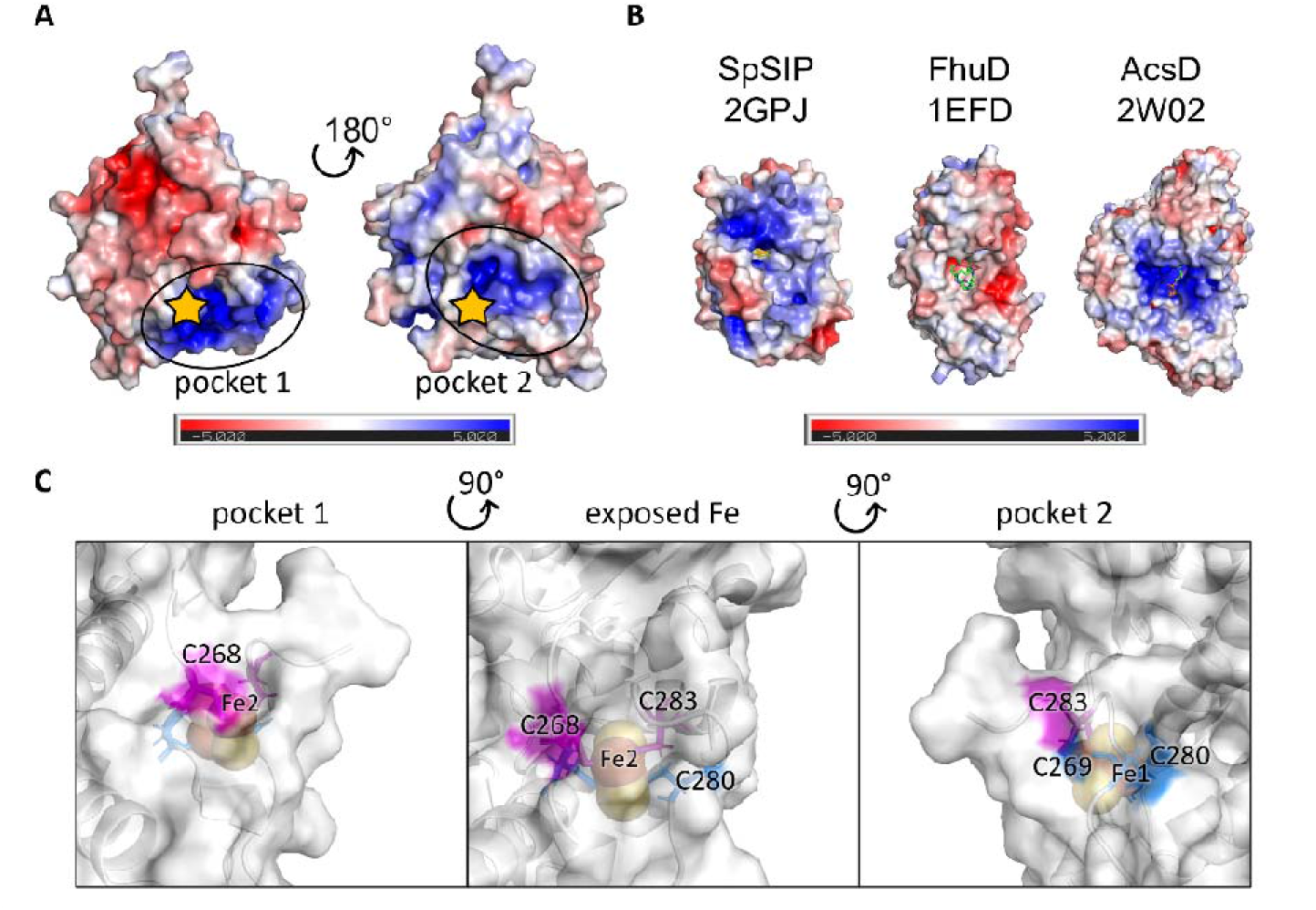
Electrostatics of FhuF and putative binding pockets. A) Electrostatic surface potential (−5 to +5 kT/e) of FhuF calculated with the APBS plugin of PyMol reveals putative binding pockets, where the star indicates the location of the 2Fe-2S cluster. B) Electrostatic surface potential (−5 to +5 kT/e) of other proteins that interact with siderophores. C) Close-up between pocket 1 and 2 showing that the 2Fe-2S cluster of FhuF sits on a ridge between these two positively charged pockets, exposing one of the Fe atoms (Fe2) to the solvent.

Interestingly, when we compare the electrostatic surface potential of FhuF with other proteins which interact with siderophores, pocket 2 of FhuF resembles the pocket found in siderophore synthetases (AcsD in Figure 6 B). Given this similarity, we used NMR spectroscopy to test the ability of FhuF to bind ATP, but no binding was observed (Data not shown) [43]. Additionally, the electrostatics of FhuF differ dramatically from those of proteins that transport siderophores, such as FhuD (Figure 6 B), which instead contain very negatively charged pockets. For proteins whose task is to carry siderophores but not reduce them, the negatively charged pocket guarantees that the reduction potential of the bound Fe(III)-siderophores is kept very low, making sure that the iron is not adventitiously reduced and released and that it is, instead, delivered to the appropriate destination.

## Conclusions

The first structure of an FSR reported in this work, the enzyme FhuF from E. coli K-12, opens new perspectives in the study of this class of proteins. The electrostatics of FhuF revealed a common feature for Fe(III)-siderophore reductases. A positively charged (proton-rich) pocket that surrounds the catalytic center guarantees the propitious environment for redox-Bohr effect, which is a common functional feature for both SIP and FSR families of enzymes. The coupled release of a proton upon electron transfer causes a drop in the local pH enhancing ferrous iron solubility upon reduction. Additionally, the 2Fe-2S cluster in FhuF sits on a ridge between two positively charged areas, suggesting that different binding sites exist for the different interacting partners, in a way that minimizes the release of “free” ferrous iron, which is highly reactive and leads to the generation of deleterious radicals [44].

The structure of FhuF shows that this is a novel 2Fe-2S protein, given its unprecedented fold and coordination of the 2Fe-2S cluster. This explains the atypical spectroscopic features of this protein, in particular with respect to the paramagnetic NMR spectra. Despite having a novel coordination mode, the cluster of FhuF shares some geometrical features with ferrochelatases, which are interestingly also involved in the handling of iron in the anabolic metabolism. However, there are no reports of paramagnetic NMR studies of ferrochelatases which would allow us to look deeper into similarities between the two, despite the abundant literature regarding this enzyme owing to its role in debilitating human diseases [45]. Surprisingly, in human ferrochelatase, the role of the 2Fe-2S cluster remains unclear. Although it is essential for the activity and stability of the protein, it is not directly involved in catalysis, i.e., the insertion of ferrous iron into protoporphyrin to form heme [46].

Additionally, the structural similarity of FhuF with siderophore synthetases also draws attention to a previously reported ferrichrome modification upon cellular uptake [47]. We could speculate that FhuF can also perform this reaction, which would reveal an additional layer to the role of FhuF in microbe-microbe interactions and how these proteins have shaped the evolution of iron uptake pathways. It is also interesting to note that FhuF showed very high rates of reduction of hydroxamate siderophores, which are not produced by E. coli strains. This catalytic activity of FhuF therefore reflects an opportunistic lifestyle of E. coli that enhances its ecological fitness, and which has been retained even in this laboratory strain. Additionally, E. coli strains express flavin-based ferric-siderophore reductases (SIP family) including YqjH, which is apparently specific for the reduction of ferric triscatecholates including hydrolyzed ferric-enterobactin complex [12]. Altogether, the specificity of these enzymes reflects the cost/benefit interplay of having a protein that on the one hand requires iron for its maturation but on the other hand can facilitate iron release very efficiently into the cellular metabolism.

The metabolic positioning of FhuF remains open for discussion. It was previously observed from pull-down assays that FhuF interacts with OppD which is annotated as an Oligopeptide transport ATP-binding protein and with YdbK annotated as a probable pyruvate-flavodoxin oxidoreductase [48]. OppD may function as a siderophore transporter of unknown substrate preference whereas YbdK is a putative electron donor for FhuF. Given the low reduction potential of FhuF, it is likely that its electron donor is a very low reduction potential donor. The pyruvate oxidation activity of YbdK was tested in vitro using Methyl Viologen as electron acceptor, which has a reduction potential in the appropriate range to make the reaction thermodynamically feasible. Furthermore, both FhuF and YdbK are upregulated under oxidative stress signaling. These conditions exacerbate the need for iron, and the reduction of FhuF by YdbK allows it to reduce ferric-siderophores, making iron available for anabolic metabolism. In this respect, it is interesting to note that the expression of FhuF is also linked to the abundance of downstream intracellular iron sinks as can be deduced by the observed up-regulation of FhuF in E. coli overexpressing ferritin [49].

This work provides for the first time a detailed structural knowledge for the comparative studies of representatives of both families of cytoplasmic ferric-siderophore reductases. This will enable further understanding of the role and mechanistic properties of these enzymes, information of crucial relevance when considering therapeutic intervention via targeting of iron uptake, which is essential for nearly all known organisms [50,51]. Indeed, within the siderophore pathway, SIPs and FSRs provide a narrow choke point between a wide diversity of cell surface siderophore receptors and intracellular iron-containing or storage proteins [52]. Inhibitors for this crucial step on the siderophore pathway can potentially reduce access to iron for anabolic metabolism and stop the recycling of apo-siderophores for subsequent rounds of iron uptake.

## Acknowledgements

The authors are grateful to Prof Alfred Trautwein, who established the first contact with Prof Berthold Matzanke, who graciously made the FhuF expression system available that allowed this work to be performed. To Kelly Frade, for the early help in finding the most stabilizing buffer for FhuF. To Paula Chicau and the N-terminal sequencing facility at ITQB for providing the measurements used in this work.

The X-ray diffraction data collection was performed at XALOC beamline at ALBA Synchrotron with the collaboration of ALBA staff. This work benefited from access to CERMAX, ITQB-NOVA, Oeiras, Portugal with equipment funded by FCT, project AAC 01/SAICT/2016. Financial support was provided by European EC Horizon2020 TIMB3 (Project 810856) and COST Action CA21115 Iron–sulphur (FeS) clusters: from chemistry to immunology (FeSImmChemNet). Financial support was also provided by Project MOSTMICRO-ITQB with refs UIDB/04612/2020 and UIDP/04612/2020, and LS4FUTURE Associated Laboratory (LA/P/0087/2020). N-terminal sequencing service was provided by the ITQB Research facilities. Fundação para a Ciência e a Tecnologia (FCT) Portugal is also acknowledged for funding through project PTDC/BIA-BQM/30176/2017, and through FCT PT-NMR PhD Program via PD/BD/135187/2017 to IBT.

## I. Supplementary Information

The structure of a novel ferredoxin – FhuF, a ferric-siderophore reductase from Escherichia coli K-12 with a novel 2Fe-2S cluster coordination

**Table S1.**
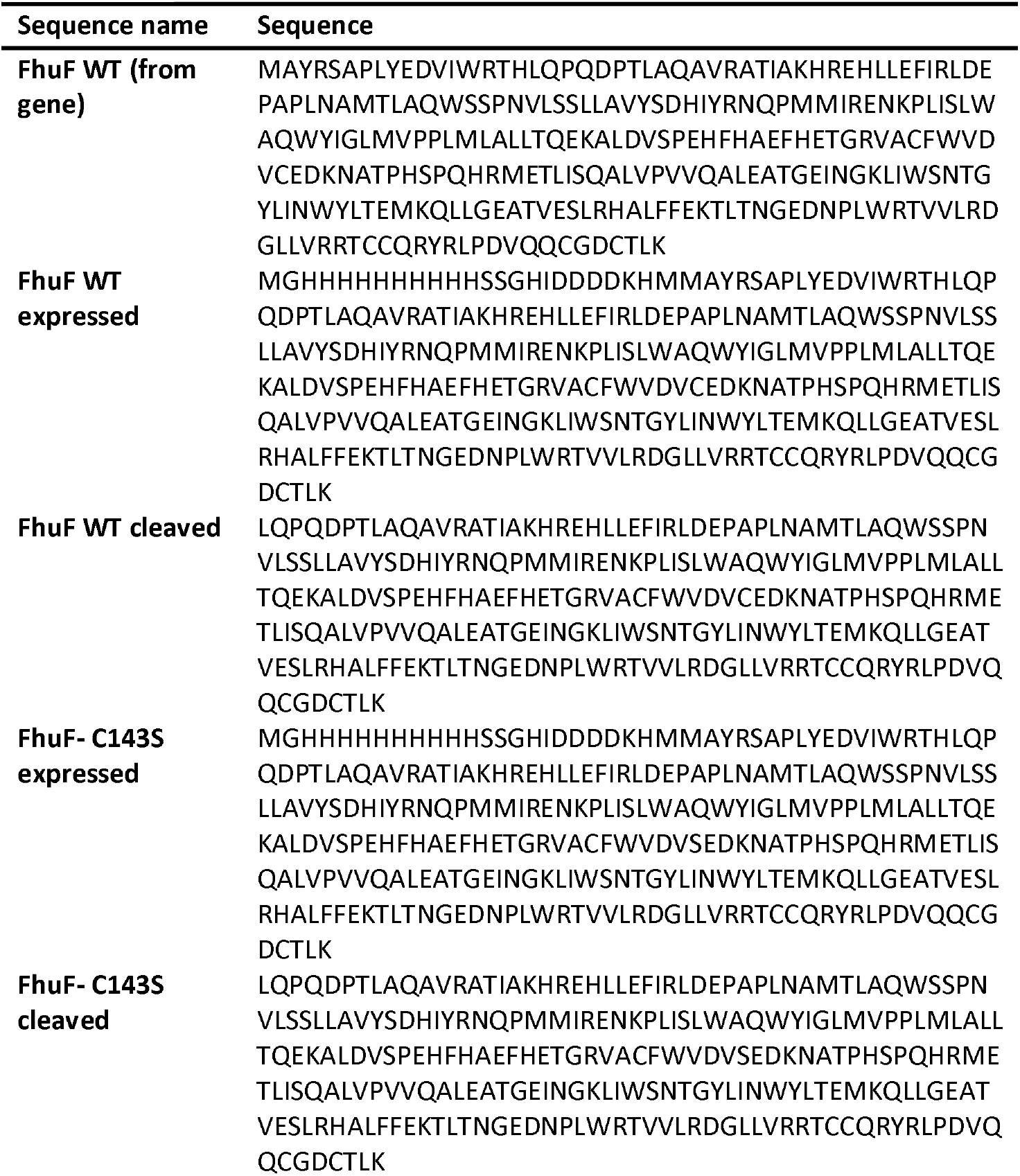
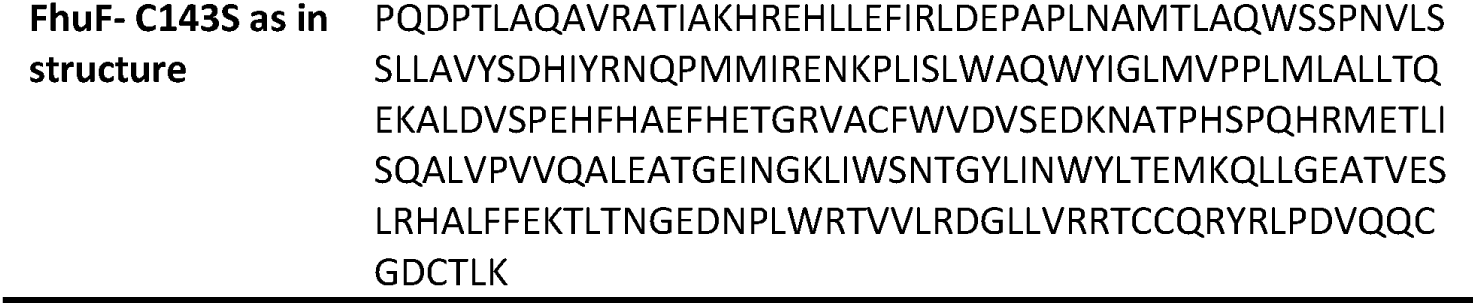
List of amino-acid sequences of FhuF obtained and reported in the manuscript.

**Figure S1.**
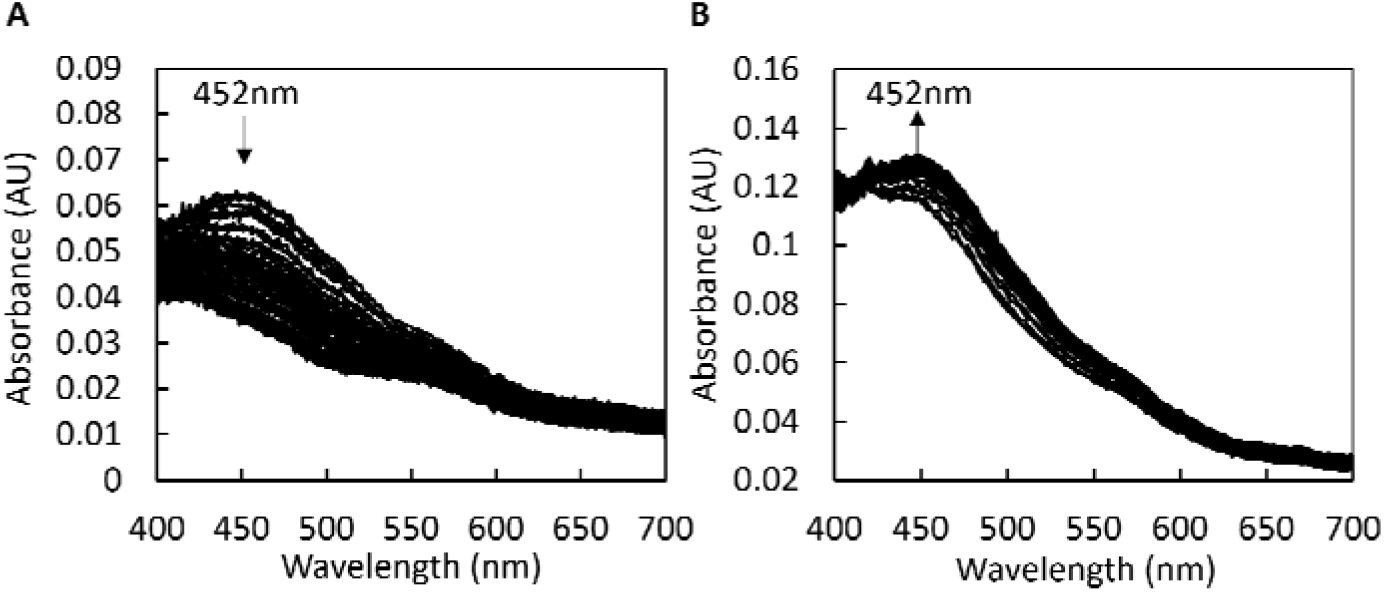
A) Representative UV-visible spectral changes upon mixing sodium dithionite with FhuF. B) Representative UV-visible spectral changes upon mixing FhuF_red_ with Fe(III)-Ferrichrome.

## References

[1] W. Martin, M.J. Russell, On the origins of cells: A hypothesis for the evolutionary transitions from abiotic geochemistry to chemoautotrophic prokaryotes, and from prokaryotes to nucleated cells, Phil. Trans. R. Soc. Lond. B. 358 (2003) 59–85.

[2] J. Liu, S. Chakraborty, P. Hosseinzadeh, Y. Yu, S. Tian, I. Petrik, A. Bhagi, Y. Lu, Metalloproteins containing cytochrome, iron-sulfur, or copper redox centers, Chem. Rev. 114 (2014) 4366–4369.

[3] R.J.P. Williams, The Bakerian Lecture, 1981 Natural Selection of the Chemical elements, Proc. R. Soc. Lond. 213 (1981) 361–397.

[4] L.B. Dugad, G.N. la Mar, L. Banci, I. Bertini, Identification of Localized Redox States in Plant-Type Two-Iron Ferredoxins Using the Nuclear Overhauser Effect, Biochemistry. 29 (1990) 2263–2271.

[5] K. Müller, B.F. Matzanke, V. Schünemann, A.X. Trautwein, K. Hantke, FhuF, an iron-regulated protein of Escherichia coli with a new type of [2Fe-2S] center, Eur J Biochem. 258 (1998) 1001–1008.

[6] M. Miethke, A.J. Pierik, F. Peuckert, A. Seubert, M.A. Marahiel, Identification and characterization of a novel-type ferric siderophore reductase from a gram-positive extremophile, J. Biol. Chem. 286 (2011) 2245–2260.

[7] I.B. Trindade, G. Hernandez, E. Lebègue, F. Barrière, T. Cordeiro, M. Piccioli, R.O. Louro, Conjuring up a ghost: structural and functional characterization of FhuF, a ferric siderophore reductase from E. coli, J. Biol. Inorg. Chem. 26 (2021) 313–326.

[8] B.F. Matzanke, S. Anemüller, V. Schünemann, A.X. Trautwein, K. Hantke, FhuF, Part of a Siderophore-Reductase System, Biochemistry. 43 (2004) 1386–1392.

[9] S.D. Springer, A. Butler, Microbial ligand coordination: Consideration of biological significance, Coord. Chem. Rev. 306 (2016) 628– 635.

[10] I. Schröder, E. Johnson, S. de Vries, Microbial ferric iron reductases, FEMS Microbiol Rev. 27 (2003) 427–447.

[11] M. Miethke, M.A. Marahiel, Siderophore-Based Iron Acquisition and Pathogen Control, Microbiol Mol Biol Rev. 71 (2007) 413–451.

[12] M. Miethke, J. Hou, M.A. Marahiel, The siderophore-interacting protein YqjH acts as a ferric reductase in different iron assimilation pathways of Escherichia coli, Biochemistry. 50 (2011) 10951–10964.

[13] K. Li, W.H. Chen, S.D. Bruner, Structure and Mechanism of the Siderophore-Interacting Protein from the Fuscachelin Gene Cluster of Thermobifida fusca, Biochemistry. 54 (2015) 3989–4000.

[14] C. Vonrhein, C. Flensburg, P. Keller, A. Sharff, O. Smart, W. Paciorek, T. Womack, G. Bricogne, Data processing and analysis with the autoPROC toolbox, Acta Crystallogr. D Biol. Crystallogr. 67 (2011) 293–302.

[15] W. Kabsch, XDS, Acta Crystallogr. D Biol. Crystallogr. 66 (2010) 125–132.

[16] A.J. McCoy, R.W. Grosse-Kunstleve, P.D. Adams, M.D. Winn, L.C. Storoni, R.J. Read, Phaser crystallographic software, J Appl Crystallogr. 40 (2007) 658–674.

[17] P. Evans, Scaling and assessment of data quality, Acta Crystallogr. D Biol. Crystallogr. 62 (2006) 72–82.

[18] P.R. Evans, G.N. Murshudov, How good are my data and what is the resolution?, Acta Crystallogr. D Biol. Crystallogr. 69 (2013) 1204–1214.

[19] J. Jumper, R. Evans, A. Pritzel, T. Green, M. Figurnov, O. Ronneberger, K. Tunyasuvunakool, R. Bates, A. Žídek, A. Potapenko, A. Bridgland, C. Meyer, S.A.A. Kohl, A.J. Ballard, A. Cowie, B. Romera-Paredes, S. Nikolov, R. Jain, J. Adler, T. Back, S. Petersen, D. Reiman, E. Clancy, M. Zielinski, M. Steinegger, M. Pacholska, T. Berghammer, S. Bodenstein, D. Silver, O. Vinyals, A.W. Senior, K. Kavukcuoglu, P. Kohli, D. Hassabis, Highly accurate protein structure prediction with AlphaFold, Nature. 596 (2021) 583–589.

[20] K. Cowtan, The Buccaneer software for automated model building. 1. Tracing protein chains, Acta Crystallogr. D Biol. Crystallogr. 62 (2006) 1002–1011.

[21] P. Emsley, B. Lohkamp, W.G. Scott, K. Cowtan, Features and development of Coot, Acta Crystallogr. D Biol. Crystallogr. 66 (2010) 486–501.

[22] G.N. Murshudov, P. Skubák, A.A. Lebedev, N.S. Pannu, R.A. Steiner, R.A. Nicholls, M.D. Winn, F. Long, A.A. Vagin, REFMAC5 for the refinement of macromolecular crystal structures, Acta Crystallogr. D Biol. Crystallogr. 67 (2011) 355–367.

[23] A. Brünger, Free R value: a novel statistical quantity for assessing the accuracy of crystal structures, Nature. 355 (1992) 472–475.

[24] V.B. Chen, W.B. Arendall, J.J. Headd, D.A. Keedy, R.M. Immormino, G.J. Kapral, L.W. Murray, J.S. Richardson, D.C. Richardson, MolProbity: All-atom structure validation for macromolecular crystallography, Acta Crystallogr. D Biol. Crystallogr. 66 (2010) 12–21.

[25] S. Velankar, C. Best, B. Beuth, PDBe: protein data bank in Europe, Nucleic Acids Res. Spec. Publ. 38 (2010) 308–317.

[26] W.L. DeLano, The PyMOL Molecular Graphics System, Version 2.3. New York, NY: Schrödinger LLC., (2020).

[27] K. Diederichs, P.A. Karplus, Improved R-factors for diffraction data analysis in macromolecular crystallography, Nature. 4 (1997) 269–276.

[28] K. Diederichs, Quantifying instrument errors in macromolecular X – Ray data sets, Acta Crystallogr. D: Struct. Biol. 66 (2010) 733–740.

[29] B.W. Matthews, Solvent Content of Protein Crystals, J. Mol. Biol. 33 (1968) 491–497.

[30] I. Trindade, J.M. Silva, B.M. Fonseca, T. Catarino, M. Fujita, P.M. Matias, E. Moe, R.O. Louro, Structure and reactivity of a siderophore-interacting protein from the marine bacterium Shewanella reveals unanticipated functional versatility, J. Biol. Chem. 294 (2018) 157–167.

[31] S.J. Wilkens, B. Xia, F. Weinhold, J.L. Markley, W.M. Westler, NMR Investigations of Clostridium pasteurianum Rubredoxin. Origin of Hyperfine ^1^H, ^2^H, ^13^C, and ^15^N NMR Chemical Shifts in Iron-Sulfur Proteins As Determined by Comparison of Experimental Data with Hybrid Density Functional Calculations, J. Am. Chem. Soc. 120 (1998) 4806–4814.

[32] B.G. Malmstrom, Rack-induced bonding in blue-copper proteins, Eur. J. Biochem. 223 (1994) 711–718.

[33] S. Todorovic, M. Teixeira, Resonance Raman spectroscopy of Fe–S proteins and their redox properties, J. Biol. Inorg. Chem. 23 (2018) 647– 661.

[34] G. Caserta, L. Zuccarello, C. Barbosa, C.M. Silveira, E. Moe, S. Katz, P. Hildebrandt, I. Zebger, S. Todorovic, Unusual structures and unknown roles of FeS clusters in metalloenzymes seen from a resonance Raman spectroscopic perspective, Coord. Chem. Rev. 452 (2022) 1–13.

[35] M. Orio, J.M. Mouesca, Variation of average g values and effective exchange coupling constants among [2Fe-2S] clusters: A density functional theory study of the impact of localization (trapping forces) versus delocalization (double-exchange) as competing factors, Inorg, Chem. 47 (2008) 5394–5416.

[36] S. Gambarelli, J.M. Mouesca, Correlation between the Magnetic g Tensors and the Local Cysteine Geometries for a Series of Reduced [2Fe-2S*] Protein Clusters. A Quantum Chemical Density Functional Theory and Structural Analysis, Inorg.Chem. 43 (2004) 1441–1451.

[37] L. Skjeldal, J.L. Markley, V.M. Coghlan, L.E. Vickery, ^1^H NMR Spectra of Vertebrate [2Fe-2S] Ferredoxins. Hyperfine Resonances Suggest Different Electron Delocalization Patterns from Plant Ferredoxins, Biochemistry. 30 (1991) 9078–9083.

[38] B. Xia, D. Jenk, D.M. LeMaster, W.M. Westler, J.L. Markley, Electron-nuclear interactions in two prototypical [2Fe-2S] proteins: Selective (chiral) deuteration and analysis of ^1^H and ^2^H NMR signals from the alpha and beta hydrogens of cysteinyl residues that ligate the iron in the active sites of human ferredoxin and Anabaena 7120 vegetative ferredoxin, Arch. Biochem. 373 (2000) 328–334.

[39] H.M. Holden, B.L. Jacobson, J.K. Hurley, G. Tollin, B.-H. Oh, L. Skjeldal, Y.K. Chae, H. Cheng, B. Xia, J.L. Markley, Structure-Function Studies of [2Fe-2S] Ferredoxins, J. Bioenerg. Biomembr. 26 (1994) 67–88.

[40] L. Banci, I. Bertini, C. Luchinat, R. Pierattelli, N. v Shokhirev, F.A. Walker, Analysis of the Temperature Dependence of the 1 H and 13 C Isotropic Shifts of Horse Heart Ferricytochrome c: Explanation of Curie and Anti-Curie Temperature Dependence and Nonlinear Pseudocontact Shifts in a Common Two-Level Framework, J Am Chem Soc. 120 (1998) 8472–8479.

[41] A. Grosdidier, V. Zoete, O. Michielin, SwissDock, a protein-small molecule docking web service based on EADock DSS, Nucleic Acids Res. Spec. Publ. 39 (2011) 1–8.

[42] A. Grosdidier, V. Zoete, O. Michielin, Fast docking using the CHARMM force field with EADock DSS, J. Comput. Chem. 32 (2011) 2149–2159.

[43] S. Schmelz, N. Kadi, S.A. McMahon, L. Song, D. Oves-Costales, M. Oke, H. Liu, K.A. Johnson, L.G. Carter, C.H. Botting, M.F. White, G.L. Challis, J.H. Naismith, AcsD catalyzes enantioselective citrate desymmetrization in siderophore biosynthesis, Nat. Chem. Biol. 5 (2009) 174–182.

[44] D. Touati, Iron and Oxidative Stress in Bacteria, Arch. Biochem. Biophys. 373 (2000) 1–6.

[45] H.D. Basavarajappa, R.S. Sulaiman, X. Qi, T. Shetty, S. Sheik Pran Babu, K.L. Sishtla, B. Lee, J. Quigley, S. Alkhairy, C.M. Briggs, K. Gupta, B. Tang, M. Shadmand, M.B. Grant, M.E. Boulton, S. Seo, T.W. Corson, Ferrochelatase is a therapeutic target for ocular neovascularization, EMBO Mol. Med. 9 (2017) 786–801.

[46] D. Pain, A. Dancis, Roles of Fe-S proteins: From cofactor synthesis to iron homeostasis to protein synthesis, Curr Opin Genet Dev. 38 (2016) 45–51.

[47] A. Hartmannt, V. Braun, Iron Transport in Escherichia coli: Uptake and Modification of Ferrichrome, J. Bacteriol. 143 (1980) 1–10.

[48] M. Arifuzzaman, M. Maeda, A. Itoh, K. Nishikata, C. Takita, R. Saito, T. Ara, K. Nakahigashi, H.C. Huang, A. Hirai, K. Tsuzuki, S. Nakamura, M. Altaf-Ul-Amin, T. Oshima, T. Baba, N. Yamamoto, T. Kawamura, T. Ioka-Nakamichi, M. Kitagawa, M. Tomita, S. Kanaya, C. Wada, H. Mori, Large-scale identification of protein-protein interaction of Escherichia coli K-12, Genome Res. 16 (2006) 686–691.

[49] H. Abdul-tehrani, A.J. Hudson, Y. Chang, A.R. Timms, C. Hawkins, J.M. Williams, P.M. Harrison, J.R. Guest, S.C. Andrews, Ferritin Mutants of Escherichia coli Are Iron Deficient and Growth Impaired, and fur Mutants are Iron Deficient, J. Bacteriol. 181 (1999) 1415–1428.

[50] S.C. Andrews, A.K. Robinson, F. Rodríguez-Quiñones, Bacterial iron homeostasis, FEMS Microbiol Rev. 27 (2003) 215–237.

[51] R.C. Hider, X. Kong, Chemistry and biology of siderophores, Nat Prod Rep. 27 (2010) 637–657.

[52] K. Honarmand Ebrahimi, P.L. Hagedoorn, W.R. Hagen, Unity in the biochemistry of the iron-storage proteins ferritin and bacterioferritin, Chem. Rev. 115 (2015) 295–326.

