## Supplementary material for "The structure of a novel ferredoxin – FhuF, a ferric-siderophore reductase from E. coli K-12 with a novel 2Fe-2S cluster coordination": Suplementary table 1

### I. Supplementary Information

**Table S1** List of amino-acid sequences of FhuF obtained and reported in the manuscript.

| Sequence name | Sequence |
|---|---|
| **FhuF WT (from gene)** | MAYRSAPLYEDVIWRTHLQPQDPTLAQAVRATIAKHREHLLEFIRLDE PAPLNAMTLAQWSSPNVLSSLLAVYSDHIYRNQPMMIRENKPLISLW AQWYIGLMVPPLMLALLTQEKALDVSPEHFHAEFHETGRVACFWVD VCEDKNATPHSPQHRMETLISQALVPVVQALEATGEINGKLIWSNTG YLINWYLTEMKQLLGEATVESLRHALFFEKTLTNGEDNPLWRTVVLRD GLLVRRTCCQRYRLPDVQQCGDCTLK |
| **FhuF WT expressed** | MGHHHHHHHHHHSSGHIDDDDKHMMAYRSAPLYEDVIWRTHLQP QDPTLAQAVRATIAKHREHLLEFIRLDEPAPLNAMTLAQWSSPNVLSS LLAVYSDHIYRNQPMMIRENKPLISLWAQWYIGLMVPPLMLALLTQE KALDVSPEHFHAEFHETGRVACFWVDVCEDKNATPHSPQHRMETLIS QALVPVVQALEATGEINGKLIWSNTGYLINWYLTEMKQLLGEATVESL RHALFFEKTLTNGEDNPLWRTVVLRDGLLVRRTCCQRYRLPDVQQCG DCTLK |
| **FhuF WT cleaved** | LQPQDPTLAQAVRATIAKHREHLLEFIRLDEPAPLNAMTLAQWSSPN VLSSLLAVYSDHIYRNQPMMIRENKPLISLWAQWYIGLMVPPLMLALL TQEKALDVSPEHFHAEFHETGRVACFWVDVCEDKNATPHSPQHRME TLISQALVPVVQALEATGEINGKLIWSNTGYLINWYLTEMKQLLGEAT VESLRHALFFEKTLTNGEDNPLWRTVVLRDGLLVRRTCCQRYRLPDVQ QCGDCTLK |
| **FhuF- C143S expressed** | MGHHHHHHHHHHSSGHIDDDDKHMMAYRSAPLYEDVIWRTHLQP QDPTLAQAVRATIAKHREHLLEFIRLDEPAPLNAMTLAQWSSPNVLSS LLAVYSDHIYRNQPMMIRENKPLISLWAQWYIGLMVPPLMLALLTQE KALDVSPEHFHAEFHETGRVACFWVDVSEDKNATPHSPQHRMETLIS QALVPVVQALEATGEINGKLIWSNTGYLINWYLTEMKQLLGEATVESL RHALFFEKTLTNGEDNPLWRTVVLRDGLLVRRTCCQRYRLPDVQQCG DCTLK |
| **FhuF- C143S cleaved** | LQPQDPTLAQAVRATIAKHREHLLEFIRLDEPAPLNAMTLAQWSSPN VLSSLLAVYSDHIYRNQPMMIRENKPLISLWAQWYIGLMVPPLMLALL TQEKALDVSPEHFHAEFHETGRVACFWVDVSEDKNATPHSPQHRME TLISQALVPVVQALEATGEINGKLIWSNTGYLINWYLTEMKQLLGEAT VESLRHALFFEKTLTNGEDNPLWRTVVLRDGLLVRRTCCQRYRLPDVQ QCGDCTLK |

| | |
|---|---|
| **FhuF- C143S as in structure** | PQDPTLAQAVRATIAKHREHLLEFIRLDEPAPLNAMTLAQWSSPNVLS SLLAVYSDHIYRNQPMMIRENKPLISLWAQWYIGLMVPPLMLALLTQ EKALDVSPEHFHAEFHETGRVACFWVDVSEDKNATPHSPQHRMETLI SQALVPVVQALEATGEINGKLIWSNTGYLINWYLTEMKQLLGEATVES LRHALFFEKTLTNGEDNPLWRTVVLRDGLLVRRTCCQRYRLPDVQQC GDCTLK |

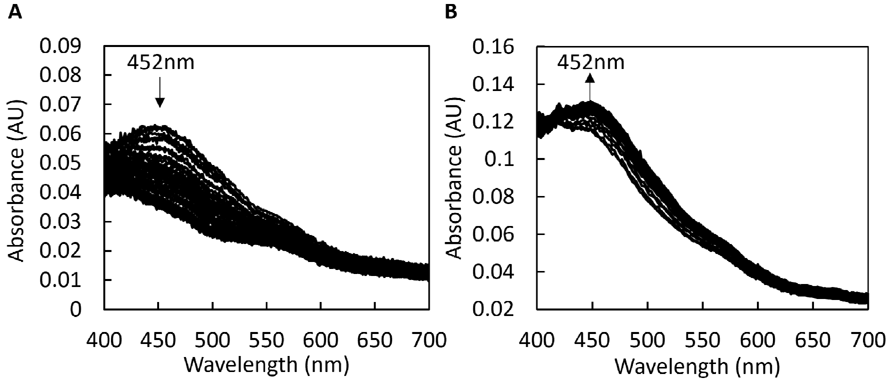


**Figure S1 A)** Representative UV-visible spectral changes upon mixing sodium dithionite with FhuF. **B)** Representative UV-visible spectral changes upon mixing $FhuF_{red}$ with Fe(III)-Ferrichrome.
